# Computational prediction of the tolerance to amino-acid deletion in green-fluorescent protein

**DOI:** 10.1101/079061

**Authors:** Eleisha L. Jackson, Stephanie J. Spielman, Claus O. Wilke

## Abstract

Proteins evolve through two primary mechanisms: substitution, where mutations alter a protein’s amino-acid sequence, and insertions and deletions (indels), where amino acids are either added to or removed from the sequence. Protein structure has been shown to influence the rate at which substitutions accumulate across sites in proteins, but whether structure similarly constrains the occurrence of indels has not been rigorously studied. Here, we investigate the extent to which structural properties known to covary with protein evolutionary rates might also predict protein tolerance to indels. Specifically, we analyze a publicly available dataset of single–amino-acid deletion mutations in enhanced green fluorescent protein (eGFP) to assess how well the functional effect of deletions can be predicted from protein structure. We find that weighted contact number (WCN), which measures how densely packed a residue is within the protein’s three-dimensional structure, provides the best single predictor for whether eGFP will tolerate a given deletion. We additionally find that using protein design to explicitly model deletions results in improved predictions of functional status when combined with other structural predictors. Our work suggests that structure plays fundamental role in constraining deletions at sites in proteins, and further that similar biophysical constraints influence both substitutions and deletions. This study therefore provides a solid foundation for future work to examine how protein structure influences tolerance of more complex indel events, such as insertions or large deletions.

## Introduction

Evolutionary change in proteins occurs via two broad classes of events: amino-acid substitutions and insertions/deletions (indels). These evolutionary events are typically subject to strong purifying selection, such that they can only occur if the resulting protein sequence can still fold and function properly. The influence of biophysical constraints on the rate of amino-acid substitution has been well-characterized, and several structural properties indicating protein tolerance to substitution events have been identified [1]. For example, residues within the protein core are less likely than are residues on the protein surface to undergo substitutions [2–5] Similarly, residues with more amino-acid contacts evolve more slowly that do sites with fewer contacts [6–10].

By contrast, the fundamental properties governing protein tolerance to indel events are comparatively understudied. Previous efforts in comparative sequence analysis have suggested that indels preferentially occur in disordered and/or loop regions and are less common in regions with secondary structure [11–15]. Furthermore, indel events may be more likely to occur on the protein surface, where residues are more exposed to solvent, rather than in a protein’s core [13, 14]. Thus, constraints on indel events appear to mirror those on substitution events, in that amino-acid substitutions are most frequent in unstructured regions and on the protein surface [1]. In spite of these observations, however, the fundamental structural properties which govern whether indel events are tolerated remain largely uncharacterized, ultimately hindering a complete understanding of how protein structure influences protein evolution.

Recently, Arpino *et al.* [16] experimentally determined the functional consequences of single deletions in enhanced green fluorescent protein (eGFP) by systematically deleting individual residues from eGFP and then assaying for function. Similar to observations from computational studies, most functional deletion mutants were located in unstructured loop regions, as opposed to in highly structured regions comprised of *β*-sheets and *α*-helices. In addition, Arpino *et al.* observed that non-functional mutants were more likely to occur in buried regions than were functional mutants [16].

Using the data from Arpino *et al.*, we investigate here how well quantities known to co-vary with amino-acid substitution rates can predict the functional consequences of deletions in eGFP. Specifically, we consider relative solvent accessibility (RSA) and weighted contact number (WCN). RSA, which ranges from 0 to 1 [17], measures how exposed a given residue is to solvent, with 0 representing a fully buried residue and 1 representing a fully exposed residue. WCN is a measure of packing density [18].

Residues with high WCN have many residue contacts and are thus tightly packed, and residues with low WCN have few contacts with other amino acids in the protein. RSA correlates positively with evolutionary rate and WCN negatively [1]. We additionally consider whether tolerance to deletions can be predicted from computational protein design [19]. Finally, we consider secondary structure (SS), which has previously been suggested as the dominant constraint on indel events [11–15].

Our results demonstrate that WCN is the best single predictor of whether a deletion will be tolerated at a given site in eGFP. RSA and protein design are good predictors, as well, but they perform somewhat worse than WCN. Interestingly, while secondary structure is a significant predictor of functional status, we find it to be less informative than WCN, RSA, and protein design. Combining multiple predictor variables yields even better performing models, with models jointly considering RSA, WCN, and protein design generally performing the best. Overall, in eGFP, the structural context of a residue appears to be a crucial factor in determining tolerance to deletion of that residue.

## Materials and Methods

### Functional data and calculation of structural properties

All functional data corresponding to each mutant was taken from Arpino *et al*. [16]. In brief, Arpino *et al*. [16] made tri-nucleotide deletions in the eGFP coding sequence using a transposon-mediated directed evolution tri-nucleotide deletion experimental approach [20, 21]. Their approach produced mutants with either a single amino-acid deletion, multiple adjacent amino-acid deletions, or an amino-acid deletion with an adjacent non-synonymous (i.e. missense) mutation. Arpino *et al*. specifically analyzed a subset of resulting mutants, categorizing each as functional or non-functional if, when expressed in *E. coli*, eGFP’s fluorescent green phenotype was preserved. In total, 87 unique mutants were assayed, of which 42 were functional and 45 were non-functional. In the following, we refer to deletion mutants as tolerated or functional if the resulting mutant fluoresced, and as non-tolerated or non-functional if the resulting mutant did not fluoresce.

Here we re-analyzed a subset of the original 87 mutants. Specifically, we identified two mutants with different nucleotide-level deletions but with the same translated product, so we only included one of these two in our analyses. We additionally excluded four non-functional mutants whose mutations had produced an early stop codon, as well as any mutants with two amino-acid changes. Finally, we excluded all mutants for which deletions had occurred in the N-terminus, C-terminus, or the eGFP chromophore. Our final dataset consisted of 72 total deletion mutants, 34 of which were functional and 38 of which were non-functional.

We computed all structural quantities using the eGFP crystal structure with PDB identifier 4EUL. We calculated the solvent-accessible surface area (ASA) for each deleted residue using DSSP [22]. To obtain relative solvent accessibility (RSA), we normalized ASA values to the maximum solvent accessibility for each amino acid in a Gly-X-Gly tripeptide, as given in Table 1 in Ref. [17]. We calculated the side-chain weighted contact number (WCN) as defined by Marcos and Echave [9]. Side-chain WCN is defined as

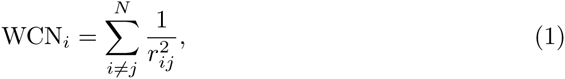

where *r*_*ij*_ is distance between the geometric center of the side-chain atoms of residue *i* and the geometric center of the side-chain atoms of residue *j* in a protein that is *N* residues long. For glycine residues, we considered the *C*_*α*_ carbon instead of the geometric center of the side chain. Although previous studies have often calculated WCN using *C*_*α*_ atoms as the reference center point, recent work has shown that using the center of mass of the entire side-chain results in stronger correlations between WCN and evolutionary variability [9, 10], and therefore we use side-chain WCN throughout this study as well. Finally, we used secondary structure (SS) as assigned previously [16].

### Modeling deletion mutations with protein design

We based all protein design work on the eGFP crystal structure with PDB identifier 4EUL. eGFP’s structure consists of a beta-barrel composed of eleven beta sheets surrounding an alpha helix which contains eGFP’s chromophore, the structural component that produces the protein’s characteristic green fluorescent phenotype. Because the chromophore interferes with commonly available protein design software, we had to first design a protein structure in which the mature chromophore was replaced with the original three amino acids (Thr65-Tyr66-Gly67) that autocatalytically form the chromophore.

To generate this pre-maturation structure, we first designed an eGFP structure with the chromophore removed, using RosettaModel [19, 23]. We subsequently used Rosetta’s relax protocol [24, 25] to optimize and re-pack the side-chains. We created 100 structures in Rosetta using the relax protocol and selected the best model as the template for design based on total score, with the best score being the most negative. We used the following commands for the relax protocol:

~~~
-database /path/to/rosetta_database
-s 4EUL_no_cro.pdb
-ignore_unrecognized_res
-use_input_sc
-constrain_relax_to_start_coords
-nstruct 100
-relax:fast
-overwrite
-out:file:scorefile relax_scorefile.fasc
-out:path:pdb ./output_pdbs/
~~~

To re-insert the three chromophore-forming residues (Thr65-Tyr66-Gly67) into the structure, we had to identify the most likely secondary structure formed by these residues. Therefore, we used Psipred [26, 27] to predict eGFP’s secondary structure elements from primary sequence. Based on the secondary structure information, we built the insert with a helical backbone, using the RosettaRemodel protocol [23]. In addition to inserting these three residues, we also designed two residues on either side of the insertion to accommodate any major structural changes created by the insertion. The following flags were used for this remodeling procedure:

~~~
-database /path/to/rosetta_database
-s 4EUL_no_cro_relaxed.pdb
-remodel:blueprint design_blueprint.txt
-run:chain A
-num_trajectory 5
-save_top 5
-ex1
-ex2
-extrachi_cutoff 1
-use_input_sc
-linmem_ig 10
-remodel:use_pose_relax
-out:file:scorefile design_protocol.fasc
-out:path:pdb ./output_pdbs/
-remodel:hb_srbb 1.0
-overwrite
~~~

We made five separate models using this protocol, and then chose the best candidate, based on overall score and visual inspection, as our wild-type template for modeling the deletion mutants.

For each of the 72 selected deletion mutants, we used Modeller [28] to create 25 initial, rough models of the deletion. We then refined these models with Rosetta, using the relax protocol with the following flags:

~~~
-database /path/to/rosetta_database
-l pdb_list.txt
-ignore_unrecognized_res
-use_input_sc
-constrain_relax_to_start_coords
-flip_HNQ
-no_optH false
-nstruct 4
-relax:fast
-overwrite
-out:file:scorefile scorefile.fasc
-out:path:pdb ./output_pdbs/
~~~

We performed four independent relaxations for each modeler structure, so that we ended up with 100 final structures corresponding to each mutant (25 Modeller models × 4 relaxed structures each). We used the mean Rosetta score for each of the 100 structures as a predictor of deletion tolerance.

### Statistical analysis of functional status

We used two different machine learning approaches, logistic regression and a support vector machine (SVM), to predict functional status using structural predictors. We employed both machine learning approaches for all models examined. We considered WCN, RSA, SS, and/or the mean score from protein design as our structural predictors. The mean score is calculated as the average Rosetta score from each of the 100 modeled structures per mutant.

We conducted logistic regressions using the `glm()` function in the statistical language R [29] with the argument `family = binomial` to specify the logistic link function. For SVM, we implemented a supervised support vector machine algorithm using the R package e1071 [30]. Our SVM used a radial basis kernel with default parameters, that is, *γ* = 1/*d*, where *d* is the dimension of the data, and the default cost of constraint violation is set to *C* = 1.

For each machine learning approach, we performed 10-fold cross-validation for each model. For each dataset, all points in the nine other datasets were used as a training dataset to train a model. The trained model was then used to predict the functional consequence (tolerated or non-tolerated) for the remaining mutants. We obtained Receiver-Operating Characteristic (ROC) curves for each model by pooling all of the predictions from the ten test datasets and plotting the true positive rate versus the false positive rate in this pooled dataset. We used the Area Under the Curve (AUC) value for each resulting ROC curve to assess the predictive power of each model. We call these AUC values the *cross-validated AUC*. Finally, for each model, we performed the 10-fold cross-validation 100 times, and we calculated the mean and standard error of the cross-validated AUC for each model.

All scripts and data from this study are freely available at https://github.com/wilkelab/eGFP_deletion_prediction.

## Results

### Variation in structural properties between non-tolerated and tolerated deletions

To examine the relationship between protein structure and tolerance to deletion, we used functional data for 72 single amino-acid eGFP deletion mutants from Arpino *et al.* [16] (see also Fig 1). We refer to a given deletion mutant as tolerated and/or functional if the mutated eGFP exhibited a fluorescent phenotype, and we refer to a given deletion mutant as non-tolerated and/or non-functional if the fluorescence phenotype was lost.

**Fig 1.**
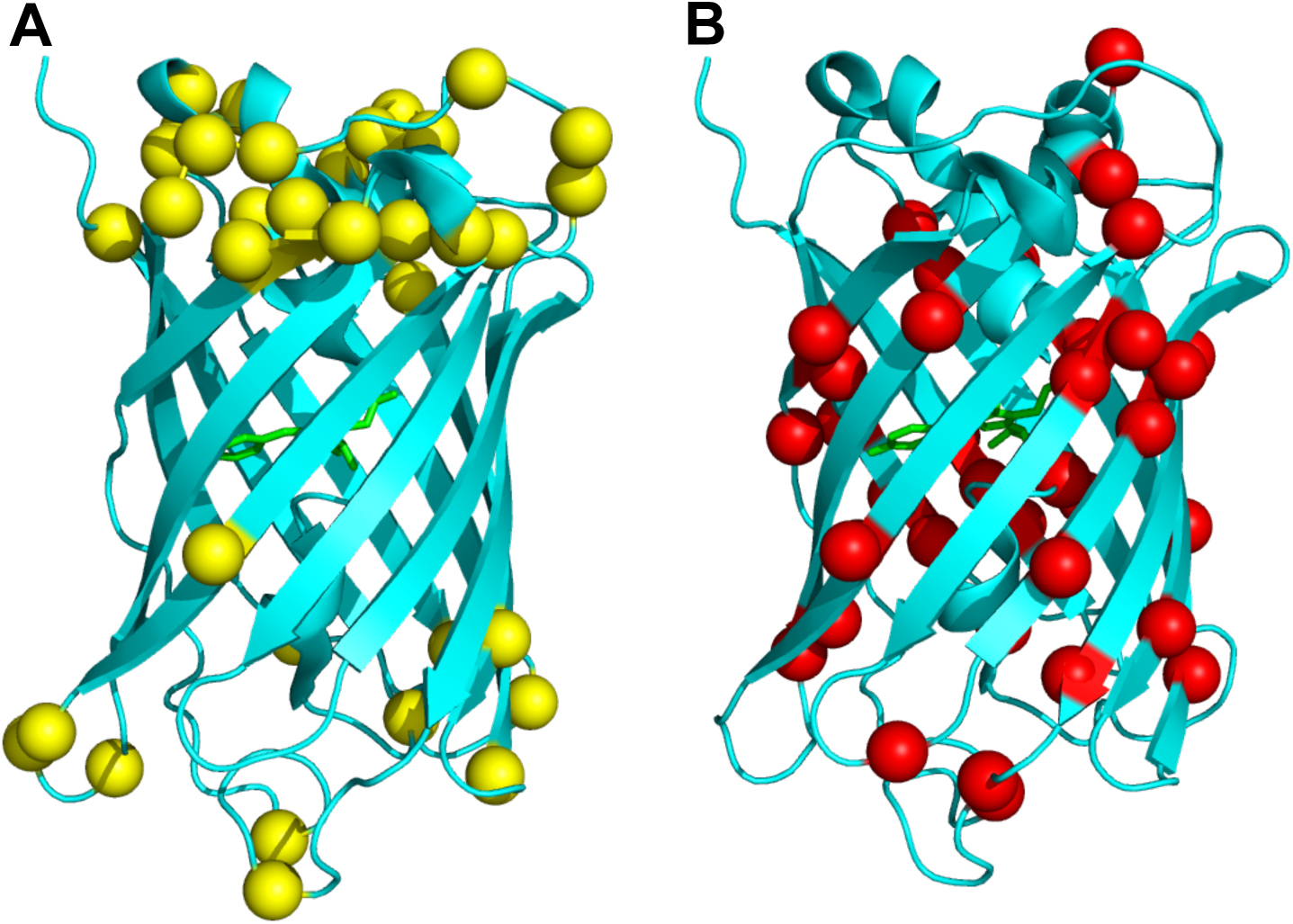
Tolerated and non-tolerated deletions in enhanced GFP (eGFP). The eGFP backbone is shown in cyan, and the chromophore responsible for florescence is shown in green. (A) Tolerated deletions. (B) Non-tolerated deletions. The C_*α*_ carbon of each deleted residue is represented by a sphere. Deletions that resulted a functioning protein (tolerated deletions) are colored yellow (A) and deletions that resulted in a non-functional protein (non-tolerated) are colored red (B). Tolerated deletions seem to be clustered towards the top and bottom of the structure, whereas non-tolerated deletions occur throughout the protein.

To characterize the structural environment of each deletion, we measured relative solvent accessibility (RSA) and weighted contact number (WCN) in the original eGFP structure for each of the 72 deleted residues. We additionally categorized each deleted residue as either beta sheet (sheet), alpha helix (helix), or loop (loop), based on the annotations previously provided [16]. We also used computational protein design to predict structures for each of the 72 mutant eGFPs. Since the eGFP structure contains a chromophore, which interferes with standard modeling approaches, we first used RosettaRemodel [19,23] to generate a model of the eGFP structure without its mature chromophore (Fig 2A). We then used a pipeline consisting of Modeller [28] and Rosetta relax [24,25] to design 100 models for each of the 72 mutant eGFPs (Fig 2B). We used the mean Rosetta score over these 100 models as a measure of how energetically (dis)favored a given deletion is, and we will refer to this score as *mean score* for the remainder of this work.

**Fig 2.**
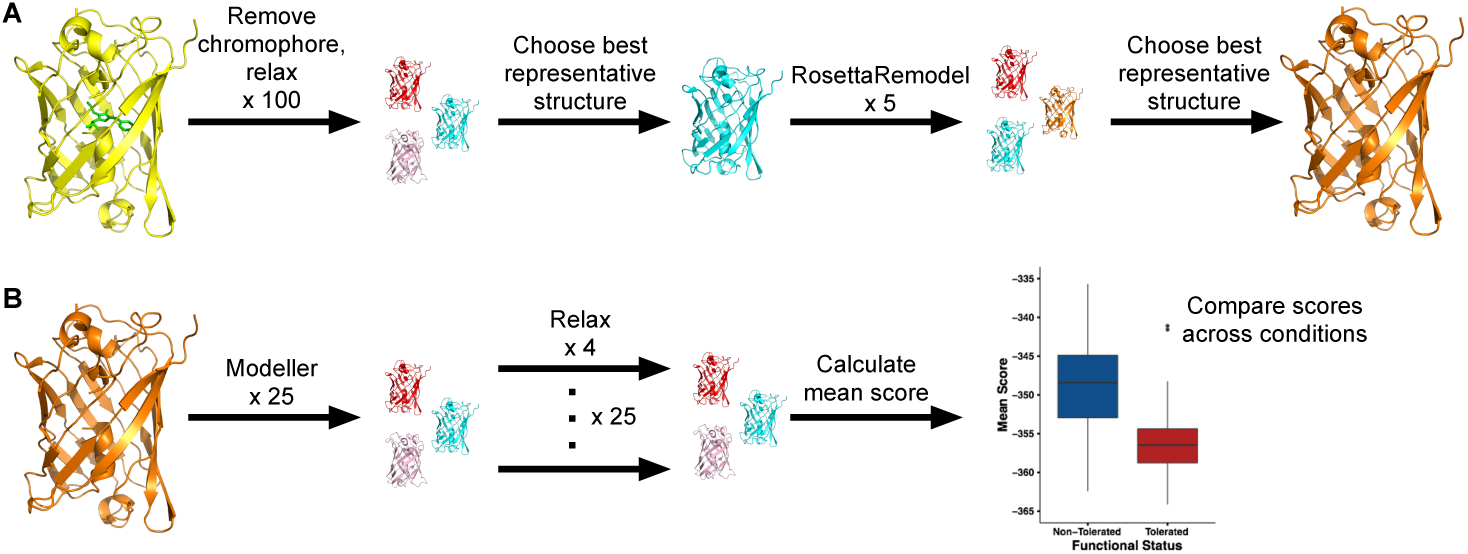
Visualization of the computational modeling pipeline. Colors represent variation between structural models produced by each protocol. (A) Generating an eGFP structure without chromophore. We first removed the chromophore and used Rosetta relax to optimize the pared-down structure. We then chose the lowest scoring model (best representative structure) from the 100 relaxed structures and reinserted the missing residues using RosettaRemodel. Finally, we picked the lowest scoring model from this protocol as the template for further analysis. (B) Generating deletion mutants. For each individual deletion mutant we considered, we first generated 25 structures using Modeller. We then refined these structures with Rosetta relax, generating four relaxes structures for each of the 25 Modeller structures. Finally, we calculated the mean score over these 100 models and used this score as a predictor of functional status for each mutant.

We first examined the distributions of each structural property between tolerated and non-tolerated deletion mutants, and we found that all structural quantities we considered displayed distinct distributions between mutant classes. In particular, deleted residues that resulted in non-functional mutants had, on average, lower RSA (Fig 3A, *t* test, *P* = 1.030 × 10^−3^) and higher WCN (Fig 3B, *t* test, *P* = 2.998 × 10^−7^) than did residues that resulted in functional mutants. Further, most functional mutants had deletions in unstructured loop regions, whereas most non-functional mutants had deletions in highly structured beta sheets (Fig 3C, *χ*^2^ contingency table, *P* = 3.642 × 10^−5^), as previously observed [16]. Finally, when comparing mean scores from protein design across mutant classes, we found that, on average, functional mutants had lower (more negative) scores (Fig 3D, *t* test, *P* = 2.084 × 10^−6^) than did non-tolerated mutants.

**Fig 3.**
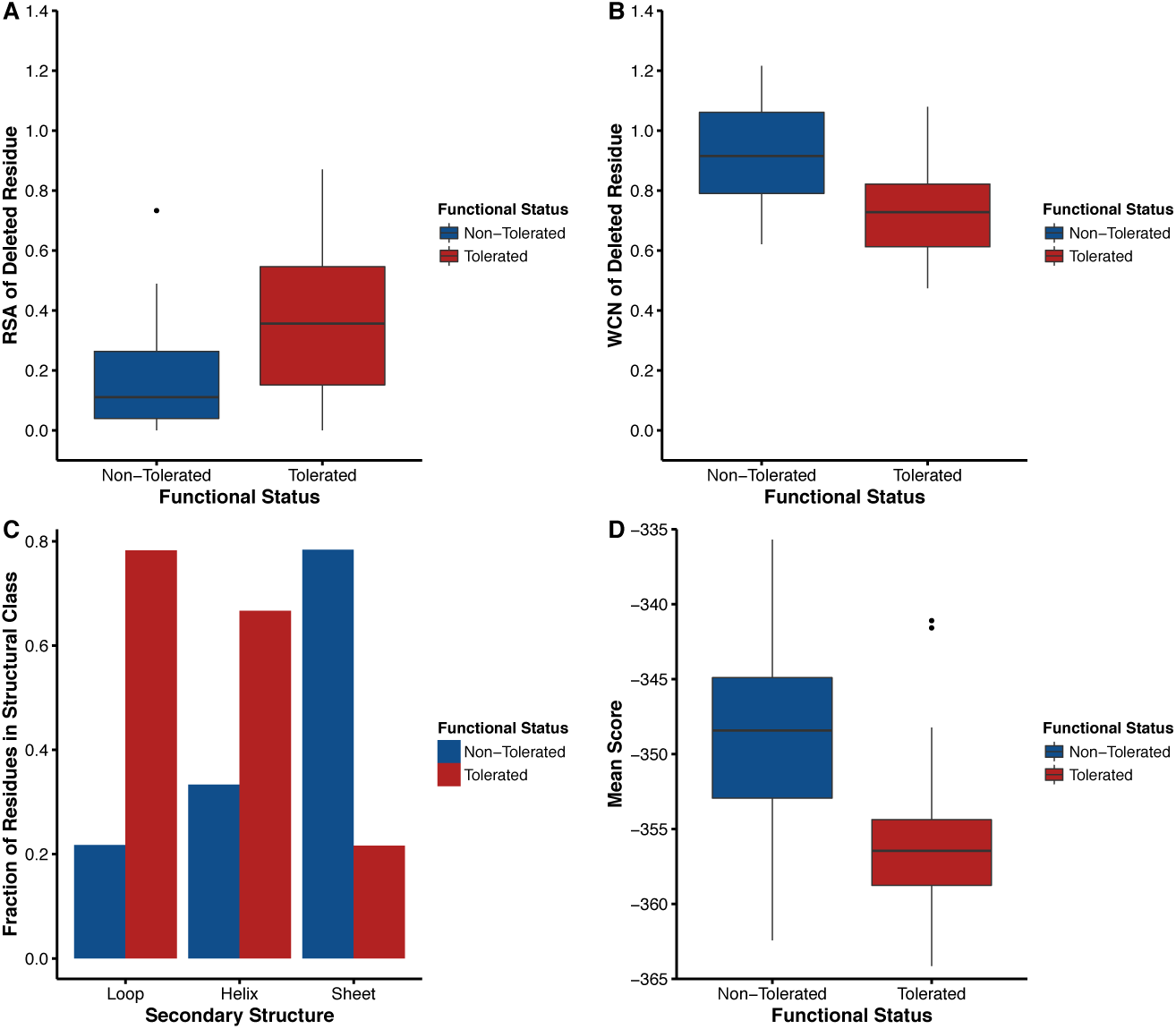
Structural properties of functional and non-functional deletions. (A) Distribution of RSA among deletions. On average, residues with tolerated deletions are more exposed than residues with non-tolerated deletions (*t* test, *P* = 1.030 × 10^−3^). (B) Distribution of WCN among deletions. On average, residues with tolerated deletions have lower WCN than residues with non-tolerated deletions (*t* test, *P* = 2.998 × 10^−7^).(C) Secondary structure of deletions. Non-tolerated deletions are colored in blue and tolerated deletions are in red. The majority of the residues deleted in the loop regions and alpha helix regions are tolerated and result in a functioning fluorescent phenotype. 78.3% and 66.7% of deleted residues are tolerated in loop and helical regions, respectively. However, only a small fraction of residues (21.6%) deleted in areas of the proteins that make up a beta sheet are tolerated. These relative frequencies are significantly different from each other (*χ*^2^ contingency table, *P* = 3.642 × 10^−5^). (D) Distribution of mean scores among deletions. Residues that are tolerant to deletion have lower scores (i.e., more negative) on average than non-tolerant residues (*t* test, *P* = 2.084 × 10^−6^).

### Prediction of functional status

The systematic variation in structural properties between tolerated and non-tolerated deletions suggested that protein structure was a viable metric for predicting tolerance to deletions. We therefore assessed how well these structural properties could directly predict the functional status of a given mutant. We constructed a series of logistic regression models using various combinations of the four structural predictors RSA, WCN, secondary structure (SS), and mean score.

For each logistic regression model, we inferred an ROC curve and calculated the Area Under the Curve (AUC) to assess the model’s predictive ability. AUC ranges from 0 to 1, where 0.5 indicates a model that performs no better than random chance and 1 indicates a model with perfect prediction ability. We additionally performed multiple rounds of 10-fold cross-validation for each model, computed an AUC value for each round of cross-validation, and recorded mean and standard error of this cross-validated AUC for each model. We used the cross-validated AUC as the primary measure of model performance.

We first used RSA and WCN individually as single predictors of functional status. Both WCN and RSA were significant predictors of deletion tolerance (Table 1), each performing better than random chance. WCN proved to be a much stronger predictor than was RSA, with a mean cross-validated AUC of 0.820 for WCN versus 0.681 for RSA.

**Table 1.**
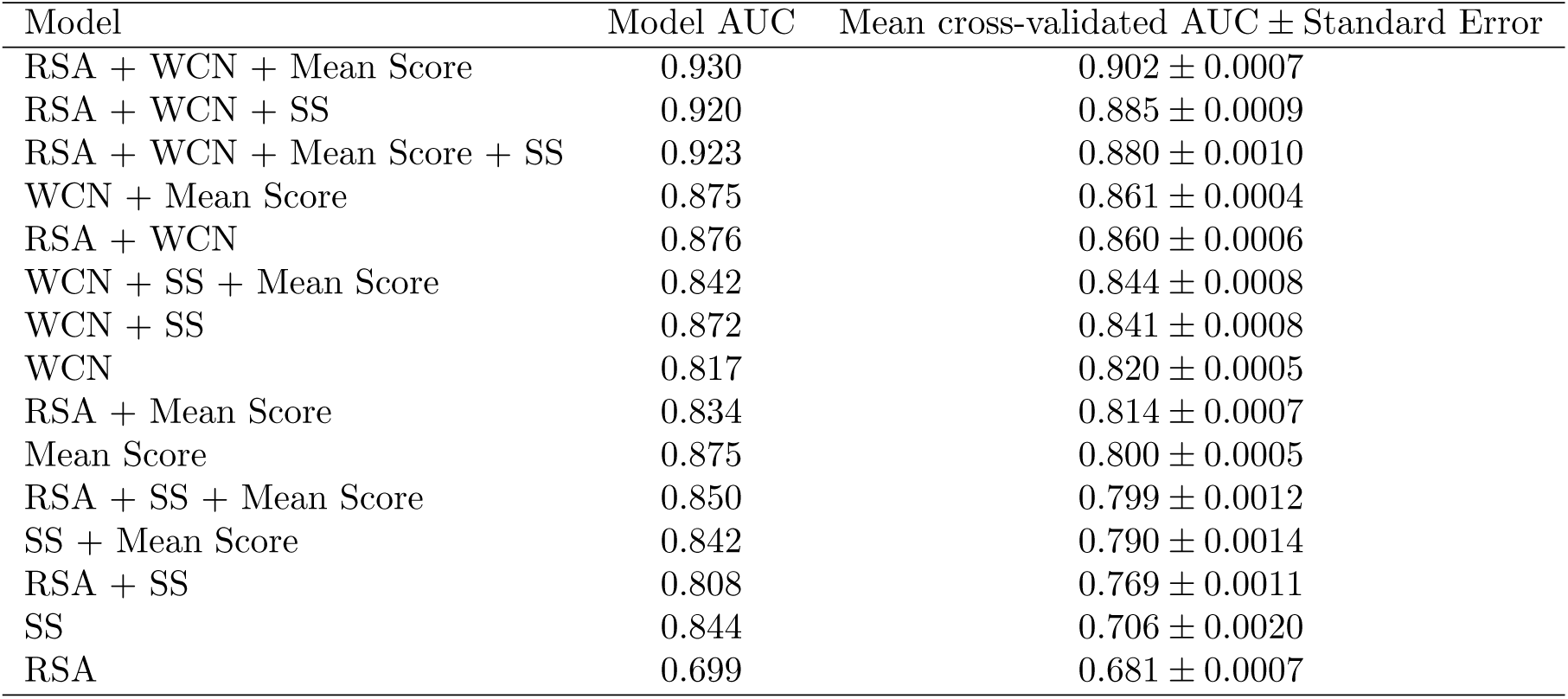
Summary of AUC values for logistic regression models. The structural properties analyzed are RSA, WCN, secondary structure (SS), and mean score. Model AUC indicates the AUC resulting from fitting a logistic regression to the entire dataset, and mean cross-validated AUC indicates the averaged AUC calculated from repeated 10-fold cross-validation. Models are sorted in order of descending cross-validated AUC.

We next included residue secondary structure (SS) classification in our logistic regressions. Secondary structure for a given residue had three possible values: beta-sheet (sheet), alpha-helix (helix), or loop (loop). The mean cross-validated AUC for a model with SS as a single predictor was 0.706, indicating that the location of residue in a given structured or unstructured region was a stronger predictor of deletion tolerance than was RSA but a weaker predictor than was WCN. Interestingly, the model AUC with SS as a single predictor was much higher than was this model’s corresponding mean cross-validated AUC value (Table 1). By contrast, model AUC and cross-validated AUC were very similar for models using WCN or RSA as predictors. This discrepancy suggest that SS models may have been more sensitive to the training set used and further that WCN and RSA were more consistent predictors compared to SS.

Finally, we used the mean score of the 100 designed structures as a single predictor of functional status. Mean score proved to be a better predictor of deletion tolerance than both RSA and SS, but WCN was still the best predictor of tolerance to deletion of a residue (Table 1). Therefore, when each predictor was considered individually, WCN emerged as the strongest predictor of functional status.

We also performed logistic regressions that incorporated these four predictors in various combinations. In general, using multiple predictors yielded better performing models, even though WCN alone beat out several combinations of other predictors, including a model using all other three predictors (RSA, SS, mean score) (Table 1). Overall, the best model for predicting functional status contained the three predictors RSA, WCN, and mean score, yielding a mean cross-validated AUC value of 0.902. This model was significantly better than the next best model, the model with RSA, WCN, and SS as predictors, which had a mean cross-validated AUC value of 0.885 (*t* test, *P* < 2.2 × 10^−16^). In fact, four of the top six logistic regression models scored by mean cross-validated AUC had mean score as a predictor, and only two had SS as a predictor. All models containing WCN as predictor performed better than did models without WCN as a predictor.

To complement our logistic regressions, we used a second machine learning approach, support vector machine (SVM) learning, to predict functional status using the same four predictors. Again, we performed 10-fold cross-validation on SVM models, and we used the mean cross-validated AUC to measure prediction accuracy. Except in the case of the model with WCN, SS, and mean score as predictors of functional status, logistic regression models yielded higher mean cross-validated AUC values compared to SVM. (Figure 4). However, differences between the two approaches were minor, and results from the SVM analysis largely agreed with those from the logistic regression analysis (Table 2). The top six scoring models were the same as those in the logistic regression analysis, although in a slightly different order (the second and third scoring models were reversed). Importantly, the model including the three predictors RSA, WCN and mean score was once again the best scoring model. With a mean cross-validated AUC value of 0.873, it was significantly better than the second best model, the model with all four predictors, that had a mean cross-validated AUC value of 0.871 (*t* test, *P* = 4.163 × 10^−6^). That including secondary structure as a predictor did not substantially improve models suggests that information contained in secondary structure may be effectively captured by more fundamental residue-level properties such as WCN and RSA.

**Table 2.** Summary of AUC values for models using a support vector machine (SVM). The structural properties analyzed are RSA, WCN, secondary structure (SS), and mean score. Model AUC indicates the AUC resulting from the entire dataset, and mean cross-validated AUC indicates the averaged AUC calculated from repeated 10-fold cross-validation. Models are sorted in order of descending cross-validated AUC.

| Model | Model AUC | Mean cross-validated AUC ± Standard Error |
| --- | --- | --- |
| RSA + WCN + Mean Score | 0.937 | 0.873 ± 0.0011 |
| RSA + WCN + Mean Score + SS | 0.937 | 0.871 ± 0.0010 |
| RSA + WCN + SS | 0.918 | 0.864 ± 0.0015 |
| WCN + SS + Mean Score | 0.901 | 0.859 ± 0.0008 |
| WCN + Mean Score | 0.908 | 0.851 ± 0.0009 |
| RSA + WCN | 0.913 | 0.845 ± 0.0014 |
| RSA + SS + Mean Score | 0.885 | 0.791 ± 0.0016 |
| WCN + SS | 0.868 | 0.788 ± 0.0016 |
| RSA + Mean Score | 0.872 | 0.787 ± 0.0020 |
| Mean Score | 0.815 | 0.767 ± 0.0014 |
| WCN | 0.820 | 0.754 ± 0.0016 |
| SS + Mean Score | 0.852 | 0.753 ± 0.0022 |
| RSA + SS | 0.821 | 0.745 ± 0.0025 |
| SS | 0.864 | 0.690 ± 0.0020 |
| RSA | 0.755 | 0.634 ± 0.0025 |

**Fig 4.**
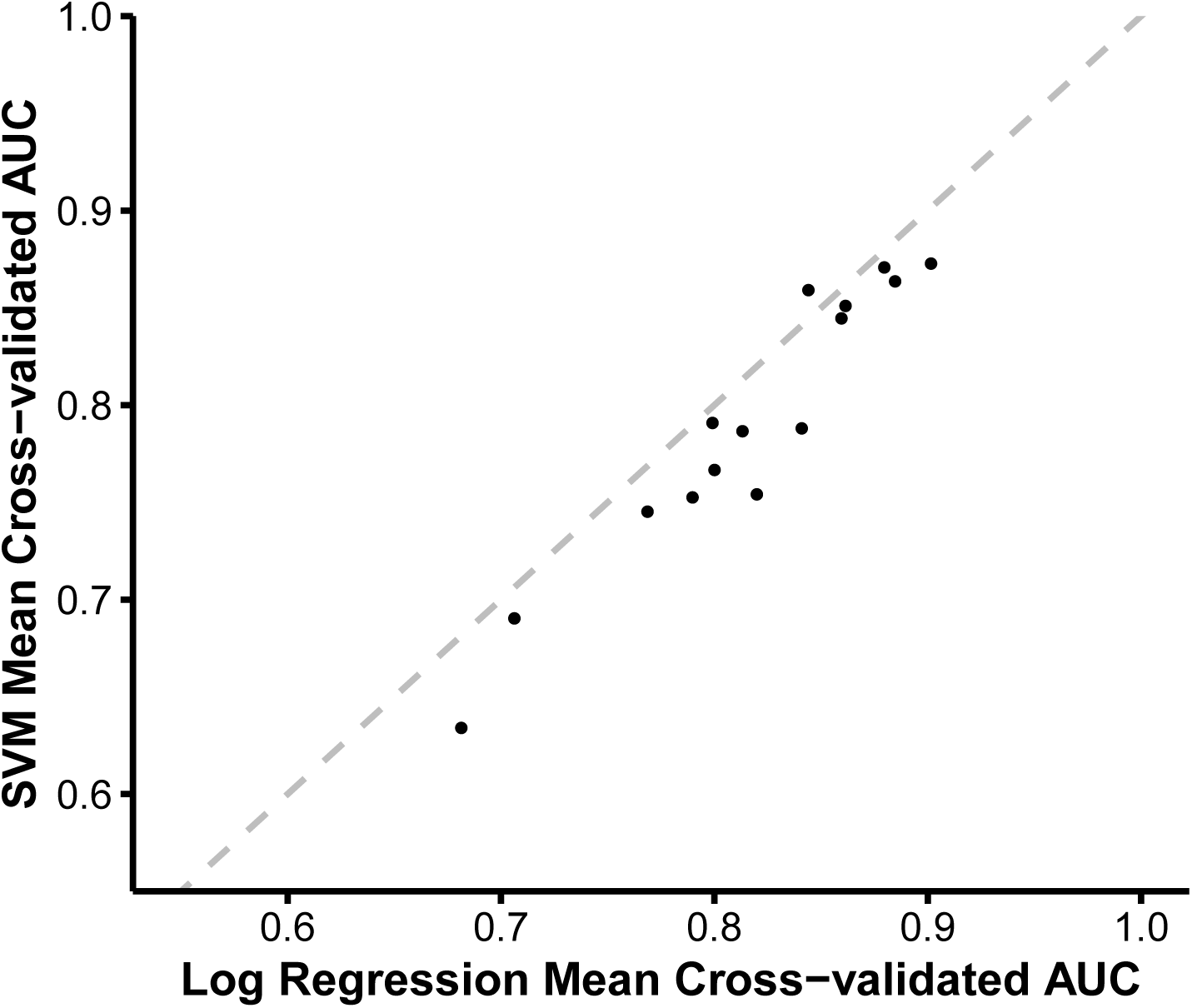
Comparison of mean cross-validated AUC from SVM and logistic regression models. For each set of predictor variables, the mean cross-validated AUC value from the corresponding SVM is plotted against the mean cross-validated AUC value from the corresponding logistic regression model. The dotted gray line represents the line *y* = *x*. For all but one model, logistic regression models with the same predictors have higher mean cross-validated AUC values.

### Covariation among structural predictors

Both logistic regression and SVM analyses revealed that including multiple predictors significantly increased prediction accuracy, but also that including all four predictors did not yield the overall best model. Moreover, secondary structure was the overall weakest predictor, a finding that seemed surprising and counter to the observation that deletions are most tolerated in loops and rarely tolerated in beta sheets (Fig 1 and Fig 3C). To gain additional insight into how the various predictors co-vary and separate tolerated from non-tolerated deletions, we performed a principal component analysis (PCA) of the structural predictor variables and visualized the response (i.e., tolerated or non-tolerated deletion) in principal component space. The PCA revealed that non-functional and functional mutants largely separated along PC1 (Fig 5A), the axis with most variance in the data. By contrast, PC2 separated sites in helices (high PC2 value) from sites in other secondary-structure regions (PC2 value near zero).

**Fig 5.**
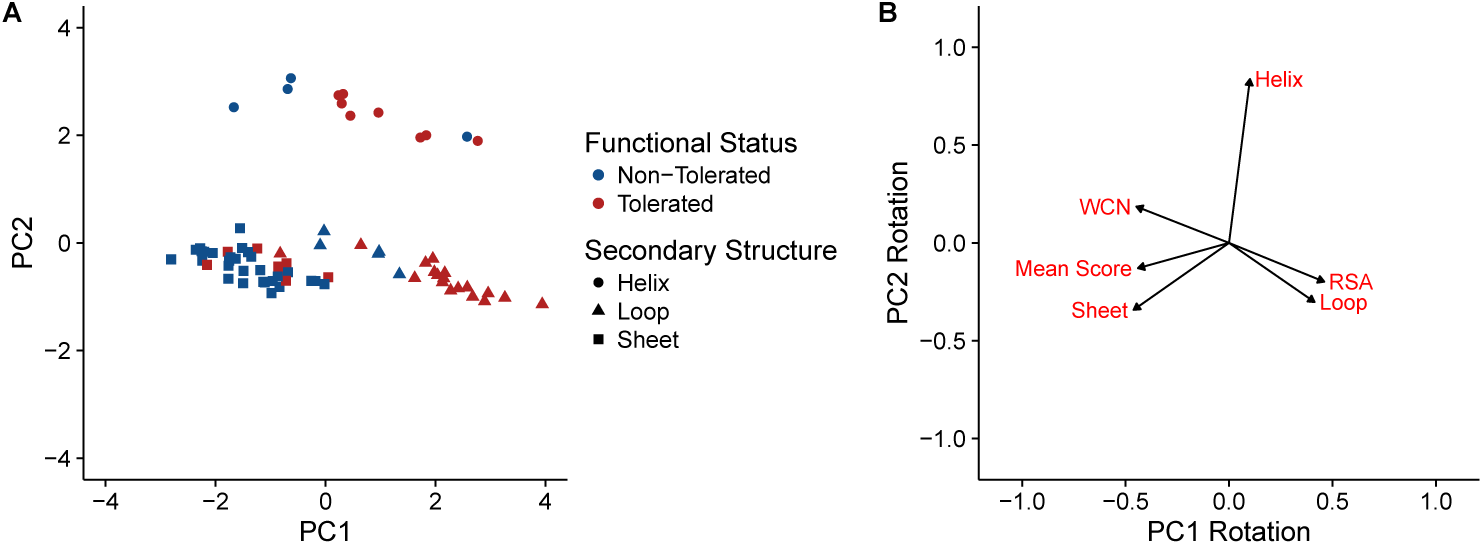
Principal Components Analysis (PCA) of the deletion dataset. (A) Deletion mutants mapped into principal components space. PC1 separates tolerated from non-tolerated mutations, while PC2 separates deletions in helices from deletions in other secondary-structure elements. (B) Visualization of the rotation matrix for the first two principal components. RSA, WCN, and secondary-structure category “Loop” are nearly collinear and primarily (but not entirely) load onto PC1. Helix is nearly orthogonal to this direction and loads primarily only PC2.

When looking at the rotation matrix (Fig 5B), we found that WCN and RSA were nearly perfectly collinear, just with opposite signs (since increasing RSA corresponds to decreasing WCN), and both were also highly collinear with the secondary structure category “Loop”. Thus, loops tended to occur primarily in regions with high RSA and low WCN, and therefore the secondary-structure classification did not provide much additional information beyond what RSA and WCN had provided. Moreover, we see from Fig 5A that not all deletions in loops were tolerated, and the non-tolerated deletions occurred at sites with lower RSA and higher WCN.

Finally, the mean score from protein design was not fully collinear with any other predictor, even though it was most similar to the secondary-structure category “Sheet”. We believe that this observation explains why mean score tended to perform better than secondary structure when added as predictor to models already containing RSA and WCN. As Fig 5A shows, not all deletions in sheets were non-tolerated, and those with lower (i.e., more negative) mean scores had a higher likelihood of being tolerated.

## Discussion

We have systematically investigated whether protein structural quantities known to predict evolutionary rate in proteins can also predict tolerance to deletions, using experimental results from enhanced green florescent protein (eGFP) as a case study. We found that models which consider the structural quantities RSA and WCN in conjunction with protein design scores provide the best predictions for whether a given deletion will be tolerated and yield a functional protein product. Overall, residues with higher RSA, lower WCN, and more negative mean scores tended to tolerate deletions more than did the converse. Our results therefore demonstrate that structural quantities known to correlate with evolutionary rate, i.e. RSA and WCN, may also be useful for predicting whether or not a protein will tolerate a deletion. As such, we find that structural considerations impose broad constraints on protein evolution, on substitutions and indels alike.

While secondary structure was a significant predictor of functional status following deletion, models with the best predictive power generally did not incorporate secondary structure. In other words, although deletions in functional mutants were typically in loop regions, and non-functional deletions occurred primarily in regions with secondary structure, secondary structure itself may not represent a strong predictor. Instead, secondary structure may in fact be a proxy for the fundamental residue-level properties, such as packing density, which actually govern protein tolerance to deletions. This finding mirrors historical trends in our understanding of biophysical constraints on amino-acid substitutions: While early work suggested that secondary structure provided the predominant constraint on evolutionary rate, advances in the field demonstrated that secondary structure was far less predictive that quantities such as WCN [1].

We further showed that protein design offers significant predictive power for a deletion’s functional status. Interestingly, this result contrasts somewhat with identified predictors of protein substitution rate, where protein design has had minimal predictive ability compared to WCN or RSA [31, 32]. For deletions, although WCN was still a better single predictor than was protein design mean score, including protein design in multi-predictor models led to consistent model improvement (Tables 1 and 2). Our principal components analysis suggested that design scores were helpful because they could distinguish sites in *α*-helices from sites in *β*-sheets, whereas WCN and RSA could not (Fig 5B). Future work will have to resolve whether this finding is specific to the beta-barrel structure of GFP or whether protein design is helpful in predicting tolerance to deletions even in structures without extensive beta sheets.

In this work, we have leveraged an existing experimental dataset [16], consisting of numerous deletions in a single protein, to delineate the biophysical constraints controlling tolerance to small deletions in proteins. Our work is markedly distinct from much of the prior work investigating where insertions and deletions (indels) can occur in proteins. Prior work investigating evolutionary pressures on indels have relied largely on sequence alignments and/or ortholog comparisons [33–36]. This approach inherently can only identify protein regions where indels are tolerated, but it cannot comprehensively reveal the evolutionary consequences of indel events. Indeed, in natural sequences, selection will have already purged non-functional proteins from the evolutionary history and only putatively functional proteins will remain. By contrast, using an experimental dataset allows us to compare directly the properties of selectively tolerated vs. non-tolerated deletion mutants.

Importantly, our study considered only the functional consequences of single–amino-acid deletion mutants, as opposed to insertions and indels spanning multiple amino-acids. Is is therefore possible that the evolutionary constraints we have identified pertain only to a small subset of indel types, and future work may elucidate how evolution constrains more complex indel events. For example, proteins appear to feature some amount of structural plasticity, allowing them to undergo compensatory structural adjustments in response to indels [13, 37–39]. It is possible that the tolerance to deletions may be specific to the compensatory abilities of a given protein, or a given protein domain.

We further emphasize that eGFP features a unique beta-barrel structure that differs from other structural families, such as globular or transmembrane proteins. Further studies will shed light on whether the constraints we have identified here pertain to proteins from different structural families. The trends we have identified in eGFP will form a robust foundation for future work to elucidate how protein structure influences the evolution of indels, ultimately providing a comprehensive picture of the forces that govern protein evolution.

## Acknowledgments

This work was supported in part by Army Research Office Grant W911NF-12-1-0390 and National Institutes of Health Grants R01 GM088344 and R01 AI120560 to COW, NIH grant F31 GM113622-01 to S.J.S., and NSF GRFP DGE-1110007 to E.L.J. The Texas Advanced Computing Center (TACC) at The University of Texas at Austin provided high-performance computing resources.

